# Long-term three-dimensional high-resolution imaging of live unlabeled small intestinal organoids using low-coherence holotomography

**DOI:** 10.1101/2023.09.16.558039

**Authors:** Mahn Jae Lee, Jaehyeok Lee, Jeongmin Ha, Geon Kim, Hye-Jin Kim, Sumin Lee, Bon-Kyoung Koo, YongKeun Park

**Affiliations:** Graduate School of Medical Science and Engineering, Korea Advanced Institute of Science and Technology (KAIST), Daejeon 34141, Republic of Korea; KAIST Institute for Health Science and Technology, Daejeon 34141, Republic of Korea; Tomocube Inc., Republic of Korea; Center for Genome Engineering, Institute for Basic Science, Daejeon 34126, Republic of Korea; Department of Physics, KAIST, Daejeon 34141, Republic of Korea; Department of Life Sciences, Pohang University of Science and Technology (POSTECH), Pohang, Republic of Korea

## Abstract

The prevailing challenges in live unlabeled high-resolution imaging of native organoids stem from technical issues like complex sample handling and optical scattering in three-dimensional architectures. In this study, we introduce low-coherence holotomography as an advanced, label-free, quantitative imaging modality, designed to overcome related technical obstacles for long-term live imaging of 3D organoids. We successfully captured high-resolution morphological intricacies and dynamic events within mouse small intestinal organoids at a subcellular resolution. Furthermore, this method provides a unique advantage in differentiating between viable and non-viable organoids, thereby expanding its potential applications in organoid-based research.

## Main

Organoids, three-dimensional (3D) multicellular structures that closely emulate organ architecture and function, are cultivated *in vitro* from both pluripotent and adult stem cells^1-3^. Organoids provide invaluable new biological tools for studying organogenesis, tissue repair, and disease pathogenesis. Although microscopy remains instrumental for organoid visualization, traditional methods, such as brightfield microscopy, fall short in capturing their 3D complexity. More advanced modalities, including confocal, multiphoton, and light-sheet microscopy, offer improved 3D imaging but come with their own caveats—namely, the necessity for fluorescent labeling of samples and potential phototoxic effects^4, 5^. Consequently, there is an ongoing quest for advanced imaging technologies that can provide high-resolution views, deep tissue penetration, and real-time observation of dynamic cellular changes of organoids, all without requiring time consuming sample preparation, labeling, and imaging, ^6^.

In recent years, quantitative phase imaging (QPI) has gained recognition as an innovative technique for the label-free imaging of live biological samples^7-10^. Within this realm, holotomography—a 3D extension of QPI—offers a unique advantage by enabling real-time capture of cellular dynamics in organoids without the drawbacks of phototoxicity or photobleaching. In this work, we employ low-coherence HT for the sustained, label-free monitoring of organoids, shedding light on their developmental trajectories and pharmacological responses. Utilizing mouse small intestinal organoids (sIOs) as our model system, we achieved time-lapse imaging over a span of 120 hours. This enabled us to monitor growth patterns, capture intricate subcellular structures with a lateral resolution of 155 nm and axial resolution of 947 nm, and quantitatively assess organoids’ drug responses. Low-coherence HT was uniquely effective in distinguishing between viable and non-viable cells within the organoids, and offered unparalleled details in depicting 3D morphological shifts following drug exposure. The method further allowed quantitative measurements of organoid volume, protein concentration, and mass, setting a new standard for comprehensive, rigorous statistical assessments for the biological studies of organoids.

In this study, the imaging protocol for mouse small intestinal organoids (sIOs) unfolds as delineated in (Fig. 1a). Initial steps involve splitting mature sIOs, and subculturing these fragments within Matrigel domes that mimic the extracellular matrix, targeting 5-10 organoids per dome. Following a five-day incubation, the organoid samples are relocated to coverslip-bottom dishes for imaging. Low-coherence HT is then employed, using four distinct patterned beams to illuminate the samples^11^. These raw images undergo a deconvolution algorithm for reconstruction. As organoids grow beyond the lateral field of view (160 µm × 160 µm) of the imaging device, tiled image capturing becomes necessary, followed by post-acquisition stitching. In cases where multiple organoids are present volumetrically, and out-of-focus beams interfere with the illumination, aberrations are inevitable. To counteract these aberration-induced artifacts, which obscure organoid boundaries, a Sobel filter is applied. This results in synthetically generated images with clearly delineated boundaries. Further segmentation and labeling of organoid outlines are accomplished using the open-source, user-interactive segmentation toolkit, ilastik^12^ (Extended Data Fig. 1). To track the axial growth of sIOs, color-coding along the axial direction is employed, with images projected in a consistent orientation. Within these segmented images, parameters such as volume, protein concentration, and mass are quantitatively assessed, enabling comprehensive, longitudinal tracking of organoid features.

**Fig 1:**
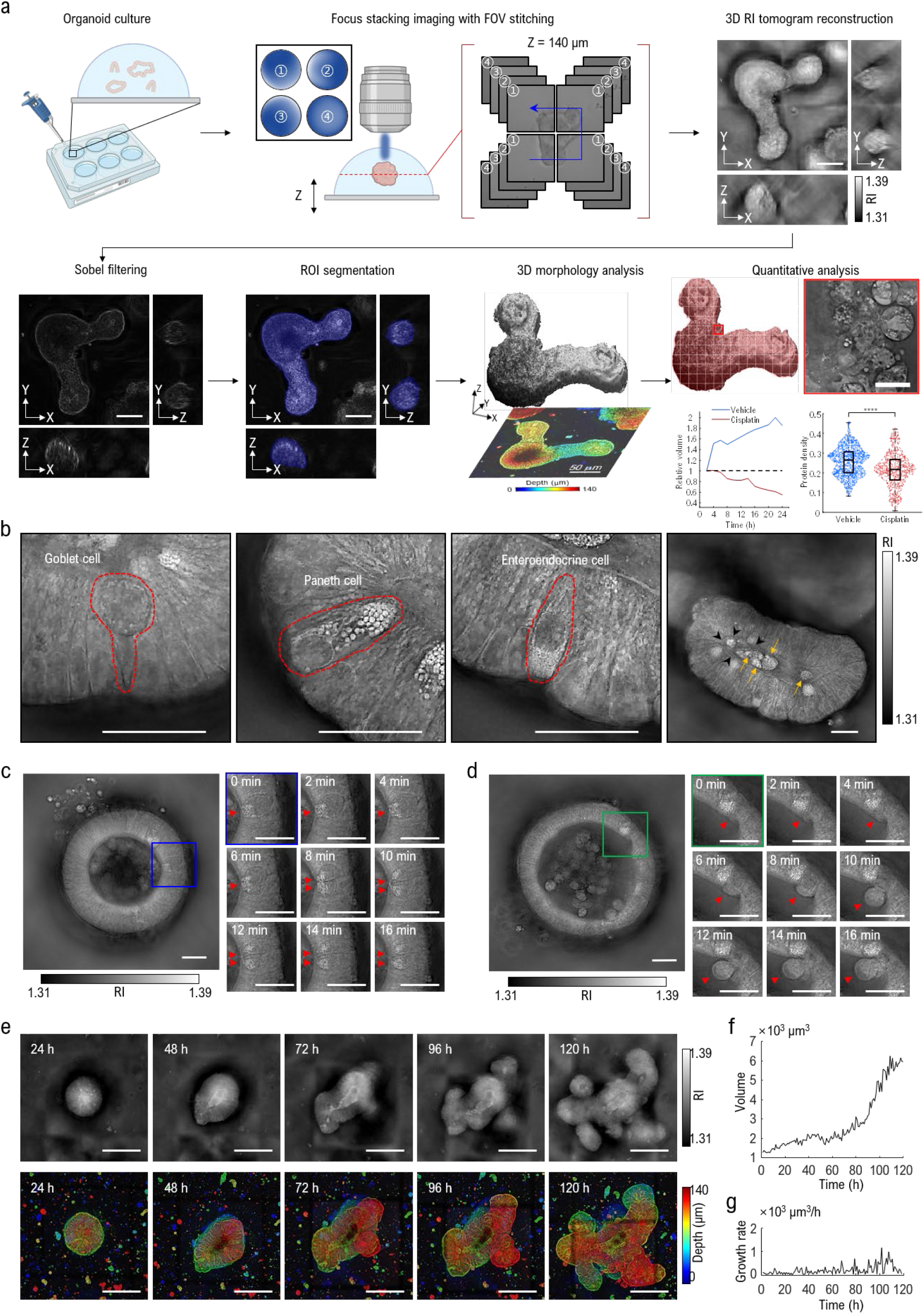
High-resolution holotomograms of live mouse sIOs via low-coherence HT. **a**, Overview of the low-coherence HT workflow. **b**, Detailed structures of sIOs. Red dotted lines represent boundaries of cells. Black arrow heads represent brush border of sIOs. Orange arrows indicate exfoliated cells and debris of dead cells. Scale bar = 20 µm. **c-d**, Time-lapse images of mitosis (c) and exfoliation (d) of cells. For each timeframe snapshot, the z-position was adjusted to the best focal plane for the targeted cells. Scale bar = 20 µm. **e**, Sequential images illustrating sIO development. For each time point, the upper row presents the focal plane, while the depth-color-coded projections are displayed in the lower row. Scale bar = 50 µm. **f-g**, A graphical illustration of sIO’s size (f) and its respective absolute growth rate (g) over time.

Through optical sectioning of reconstructed HT images, we elucidate the complex internal architecture of sIOs (Fig. 1b). Remarkably, we were able to identify goblet, Paneth, and enteroendocrine cells, each exhibiting typical morphologies consistent with previously reported electron microscopic studies^13^. Additionally, observed cellular polarity provides valuable insights into the apical lumen and basal environments. Wide-field stitched images further reveal macroscopic features such as brush border formation and the accumulation of exfoliated cells within the apical lumen.

Benefiting from the rapid acquisition capabilities of low-coherence HT, we were able to capture the dynamic behavior of live sIOs embedded in Matrigel. While specific dividing cell types were not identified, key mitotic events were observed, including apical cytokinesis and cell interspersion, followed by basal reattachment (Fig. 1c and Movie 1 online). These observations correlate well with established mitotic features from prior studies^14^. Moreover, we documented epithelial cell exfoliation (Fig 1d) and subsequent cellular apoptosis and chromatin condensation (Movie 2 online). While previous studies were limited to observing intestinal cell exfoliation^15, 16^, our study expanded upon this by capturing the entire process, including intercellular migration and apoptosis of exfoliated cells. This supports the long-debated notion that the exfoliation removes senescent or damaged enterocytes^17^.

To systematically examine sIO development, Matrigel-embedded organoids were subjected to hourly imaging over a span of 120 hours. From the initial enterocystic stage, we noted phenomena such as symmetry breaking and protrusion, eventually leading to the budding of crypts (Fig. 1e). The axial progression of cyst formation and crypt budding was vividly captured using projected depth-color-coded images. By applying segmented masks to Sobel-filtered images, we tracked organoid size over this period, revealing an exponential growth trend (Fig. 1f). Variability in organoid dimensions was noted, attributed to the shedding of apoptotic cells; these fluctuations were quantified as alterations in the absolute growth rate (Fig. 1g). Importantly, a temporal increase in growth rate was observed as the experiment progressed.

To evaluate cell death within the organoids, sIOs were treated with cisplatin, and morphological changes were closely observed. We cultured the sIOs in media containing varying concentrations of cisplatin to ascertain the optimal concentration for discernible toxic effects. A concentration of 10 μM was identified as optimal, a finding corroborated using bright-field microscopy (Extended Data Fig. 2). Dimethyl sulfoxide (DMSO) served as a vehicle and thus a positive control. Post-treatment, the samples were placed in the imaging system’s incubator and were imaged at 10-minute intervals over a 48-hour span (Fig 2a). Post-cisplatin treatment, we observed that the branched crypts underwent noticeable shrinkage and displayed an increasing number of dissociated dead cells. In contrast, the vehicle-treated organoids showed enhanced crypt budding and growth (Fig. 2b). The observed reduction in organoid volume can likely be attributed to cellular shrinkage and the exfoliation and dissociation of dead cells, as previously reported^18^. Quantitative tracking of organoid size further supported these observations. While the volume of the vehicle-treated sIOs increased, that of the cisplatin-treated sIOs decreased (Fig. 2c). Using the linearly proportional relationship between drymass density and the refractive index (RI)^19, 20^, we calculated the protein density in sIOs. The vehicle-treated group maintained a consistent protein density, whereas a drastic decrease was observed in the cisplatin-treated group after 12 hours (Fig 2c). By computing the product of organoid size and protein density, we determined the overall protein mass. Here, an increase was noted in the vehicle-treated group, while a decrease was evident in the cisplatin-treated group.

**Fig 2:**
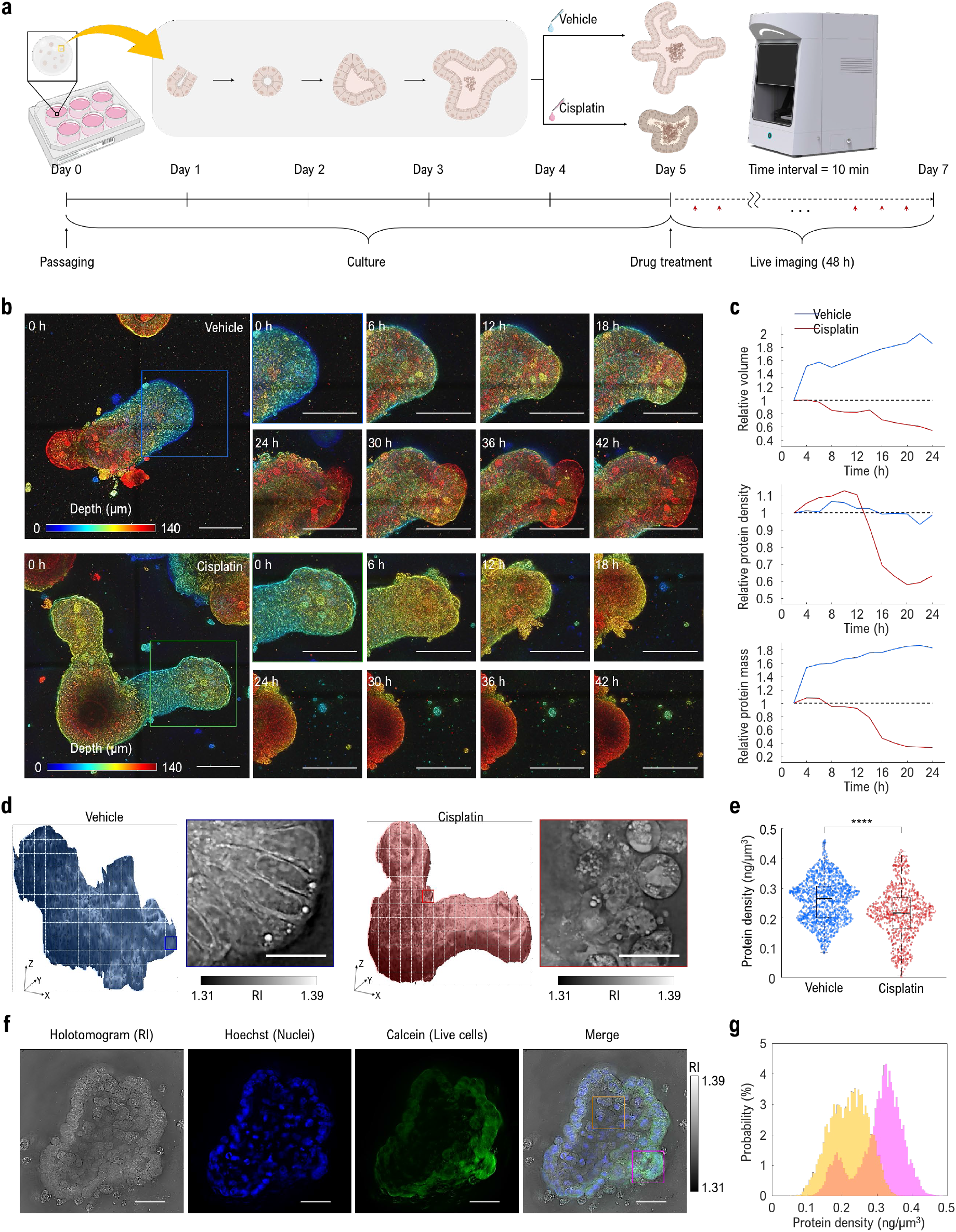
In-depth quantitative analysis of sIOs leveraging low-coherence HT. **a**, Diagram illustrating holotomogram-based viability assessment of drug-treated sIOs. Red arrowheads highlight the time points at which images were captured. **b**, Time-lapse images of sIOs following drug treatment. Each colored box is magnified, with the color bar indicating depth of samples. Scale bar = 20 µm. **c**, Longitudinal quantitative assessment of sIOs. Every parameter is calibrated against the initial measurement; the dotted line signifies the baseline value. **d**, Illustration of the subvolume sectioning strategy alongside representative images. Scale bar = 20 µm. **e**, scattered plot of protein density between vehicle-treated sIO and cisplatin-treated sIO. each dot represents a subvolume of sectioned images. **f**, Correlative images of sIO labeled with Hoechst (blue) and Calcein-AM (green). In the merged image, the magenta and orange boxes indicate live cell-dominant and dead cell-dominant regions, respectively. Scale bar = 50 µm. **g**, Histogram illustrating the protein density distribution within the boxed regions of (f). Each color corresponds to the respective color of the boxes.

To further elucidate the marked decline in protein density observed in the cisplatin-treated group, we investigated the RI distribution within the sIOs. Initially, our aim was to perform a cell-by-cell analysis by segmenting individual cells using single-cell masks. However, this approach was impeded by indistinct cell boundaries and cell clumping. To circumvent this challenge, we partitioned the holotomograms of the sIOs into multiple cubic sections, each measuring 50 μm × 50 μm laterally and 10 μm axially. This strategy ensured that each section contained an entire single enterocyte for more accurate analysis (Fig. 2d). We calculated protein density of each cubic section, and compared between organoid of cisplatin-treated sIO and vehicle-treated sIO after 24 hours from treatment (Fig 2e). The cisplatin-treated group showed lower protein density (mean ± std; 0.232 ± 0.103 ng/μm^3^), while vehicle-treated group showed higher (0.277 ± 0.085 ng/μm^3^). We hypothesized the lower protein density originate from the increased number of dead cells within the cisplatin-treated sIO since the region including dead cells exhibited low RI (Extended Data Fig 3). Previous research reported that, during cell death, a decrease in the phase of scattered beams from cells, which also indicates a decrease in RI and this back up our hypothesis^21^. To validate our hypothesis, we stained sIO with Hoechst and Calcein-AM and take their fluorescent images with low-coherence HT (Fig 2f). Since Hoechst stain whole nucleus of cells in any live/dead state while Calcein-AM can stain only live cells, we could identify live cell dominant region and dead cell dominant region. Then we compared protein density of co-stained region (live cells dominant) and Hoechst-only stained region (dead cells dominant). The histogram of protein density revealed different distribution between co-stained region and Hoechst-only region (Fig 2g). The dead cell dominant region showed lower protein density (0.220 ± 0.054 ng/μm^3^) compared to live cell dominant region (mean ± std; 0.317 ± 0.067 ng/μm^3^).

In summary, low-coherence HT offers a real-time, label-free method for organoid imaging, enabling researchers to explore the detailed complexities of organoids without time-consuming preparation and labeling. Using low-coherence HT, we captured long-term, real-time observations of biological processes within organoids, such as cell apoptosis, migration, mitosis, and dynamics of subcellular organelles within resolutions between bright field microscopy and electron microscopy. These high-resolution 3D observations were difficult to achieve in real-time using conventional imaging methods, particularly in intact organoids in Matrigel media. This study presents advancements in two primary domains. Firstly, the necessity for sample preparation steps, including fixation and staining, is obviated while preserving the structural integrity of live organoids. Secondly, the quantitative analytical capabilities of low-coherence HT provide robust various measures to assess organoid biology, including the RI values, making our imaging system a new tool for organoids in pharmacological screening applications.

## Methods

### Organoid culture

Mouse sIOs, derived from primary crypts isolated from the mouse small intestine, were kindly provided by the Center for Genome Engineering (Institute for Basic Science, Korea). Organoid culture media was either manually prepared or purchased. For sIO culture media preparation, advanced DMEM/F12 (Gibco, USA) was supplemented with 0.1M HEPES (Gibco, USA), 1X Glutamax (2mM L-alanyl-L-glutamine dipeptide) (Gibco, USA), 100U/mL Penicillin-Streptomycin (Gibco, USA), 1X serum-free B27 supplement (Gibco, USA), 1.25mM N-Acetyl-L-cysteine (NAc) (Sigma-Aldrich, USA), 0.05 μg/mL mouse EGF (Gibco), 0.1 μg/mL human Noggin (Peprotech, USA), and 0.1 μg/mL human R-Spondin-1 (Peprotech, USA). Alternatively, the commercially available culture medium, IntestiCult (STEMCELL, USA), was prepared according to the manufacturer’s instructions. The culture medium was replaced every 2-3 days, and organoids were passaged weekly. For passaging, both the Matrigel dome and the organoids were mechanically dissociated into individual crypt domains. These individual crypts were then mixed with fresh Matrigel at a split ratio of 1:8, kept at temperatures below 4°C, and distributed in 15 μL dome shapes on a 48-well plate. To polymerize the Matrigel dome, the plate was placed upside-down in an incubator for 10 minutes at 37°C with 5% CO_2_. Once the Matrigel solidified, 250 μL of fresh culture medium was added to each well.

### Imaging platform

Holotomograms of sIOs were obtained using a low-coherence HT system (HT-X1, Tomocube Inc., Korea). HT-X1 employs incoherent 450-nm LED light for illumination, enabling the imaging of the RI of thick specimens with reducing speckle noise. The sample undergoes axial scanning over a range of 140 μm, with the transmitted intensity for each illumination pattern being measured at intervals of 947 nm. Subsequently, deconvolution and stitching techniques are utilized to reconstruct the RI of the samples within the desired field of view^11^. Low-coherence HT has a theoretical resolution of 155 nm and 947 nm in the lateral and axial directions, respectively. An integrated stage top incubation chamber ensured stable physiological conditions of temperature, humidity, and CO_2_ concentration during long-term imaging over several weeks. All motorized microscopic operations were controlled and monitored by an operating software TomoStudio X (Tomocube Inc., Korea).

### Sample preparation for imaging

The HT-X1 is adaptable with a wide variety of commercial imaging dishes, multi-well plates, and custom imaging vessels, as long as they have a bottom thickness of #1.5H. In this study, organoids were imaged using a coverslip-bottomed imaging dish (Tomocube Inc., Korea). Prior to imaging, the sIOs underwent mechanical dissociation as part of the organoid passaging procedure. The dissociated individual crypts were mixed with fresh Matrigel, then distributed as 15 μL domes onto the coverslip-bottom imaging dish. This was followed by a polymerization process. Once the Matrigel domes solidified, they were topped up with 3mL of culture medium and the imaging dish was positioned in the chamber of the HT-X1.

For correlative organoid imaging, a staining solution is prepared using Calcein-AM (ThermoFisher, USA) diluted 1:200 to achieve a 5 μM concentration, and Hoechst (ThermoFisher, USA) diluted 1:1000 to reach a 1 μg/mL concentration. Both are in 3 mL of DPBS (Sigma Aldrich, USA). Following this, the culture medium surrounding the sIOs in the imaging dish is gently removed. Subsequently, sIOs are rinsed with DPBS. Once the rinse is removed, the staining solution is applied to the sIOs, ensuring uniform coverage. The organoids are then incubated for 1.5 hours to ensure optimal staining. After this duration, the excess staining solution is gently aspirated, followed by a final rinse with DPBS to remove any residual dye.

### Data acquisition

The center of the Matrigel dome was found manually with the help of a brightfield preview image scanned over the area of 4 mm × 4 mm. Randomly selected organoids within the preview were imaged with a field of view (FOV) of 160 μm × 160 μm and a depth range up to 140 μm. For the organoids exceed the size of FOV, multiple holotomograms were acquired to construct a stitched volume. The stitching process was automatically done by the software TomoStudio X (Tomocube Inc, Korea). In the case of time-lapse imaging, the humidity was maintained by replenishing the incubation chamber with deionized H_2_O 5 mL/day. Temperature and CO_2_ concentration were automatically adjusted to 37°C and 5% respectively by the chamber controller unit.

### Image processing

Each stack of raw holotomograms was processed with Matlab (Mathwork, USA). First, we calculated gradient image from raw image by applying Sobel filter for each XY-slice. Stacks were cropped with a range along z-axis determined by thresholding mean intensity of gradient, representing the existence of in-focus features within each slice. Generated stack of gradient image then applied by depth color coding. Finally, each stack was projected in single plane by maximum intensity-based projection method. Applied colormap was expressed as a color bar in each produced image.

### Feature extractions

After obtaining the organoid mask using ilastik, it was processed in Matlab to determine the volume through the built-in function ‘regionprops3’. The number of voxels was converted to volume by adjusting for both lateral and axial resolution. Protein density was derived from the organoid’s RI, with an increment value of 0.185 ng/μm used in this study. The protein mass of the organoids was then calculated by multiplying the volume by the protein density. Histograms of the organoids were generated with the Matlab built-in ‘histogram’ function. The region of interest for the histogram was specifically designated using the organoid mask to exclude non-organoid regions.

## Contributions

M.J.L. and J.H.L designed the study with input from Y.K.P. and B.K.K. Data acquisition was done by M.J.L. and J.H.L. J.H.L. performed data processing and M.J.L. performed data analysis. M.J.L. and J.H.L. wrote the manuscript, with feedback from G.K., H.J.K., S.M.L., H.H., J.M.H., B.K.K and Y.K.P.

## Acknowledgement

We thank all the members of the BMOL in KAIST, Center for Genome Engineering in IBS and Tomocube Inc. This work was supported by Basic Science Research Program through the National Research Foundation of Korea (NRF) funded by the Ministry of Education (RS-2023-00241278), NRF of Korea (2015R1A3A2066550, 2022M3H4A1A02074314), Institute of Information & Communications Technology Planning & Evaluation (IITP; 2021-0-00745) grant funded by the Korea government (MSIT), KAIST Institute of Technology Value Creation, Industry Liaison Center (G-CORE Project) grant funded by MSIT (N11230131) and Tomocube Inc. The authors thank Herve Hugonnet for technical discussion.

## Data availability

Data underlying the results presented in this paper are not publicly available at this time but may be obtained from the authors upon reasonable request.

## Extended Data

**Extended Data Figure 1:**
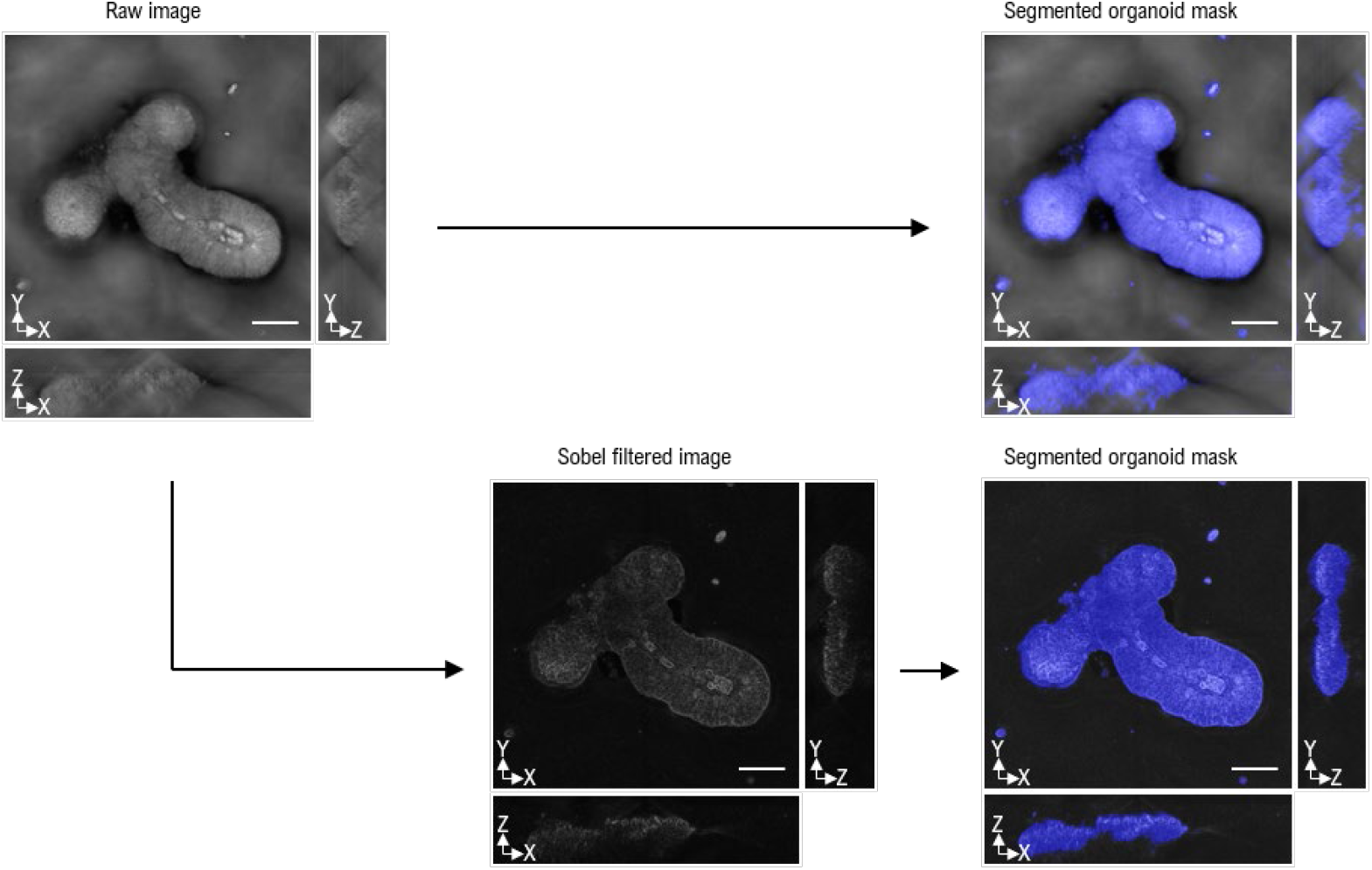
Sobel-filtered image for mask segmentation. To segment the organoid mask, a Sobel filter was applied to the raw holotomogram. Inevitable aberrations hinder image quality and blur the organoid boundaries; however, the gradient of the holotomogram clearly reveals these boundaries. Subsequently, the segmentation program, ilastik, was applied to the Sobel-filtered image of the organoids (colored in blue). It should be noted that mask segmentation using raw holotomograms yields inaccurate organoid regions, especially axially due to aberration. Scale bar = 20 µm.

**Extended Data Figure 2:**
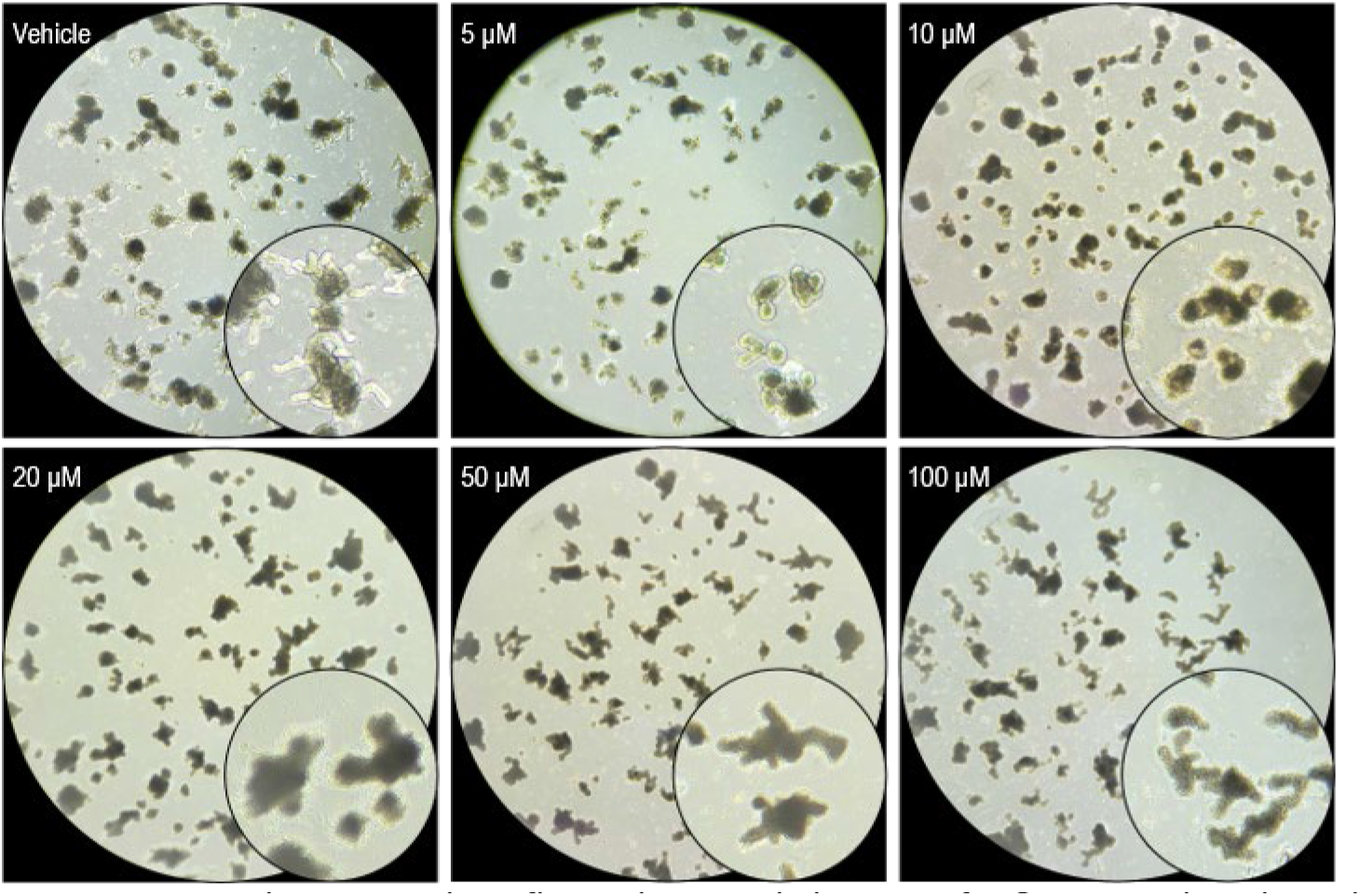
Bright-field microscopic images of sIOs at varying cisplatin concentration. To determine the optimal concentration of cisplatin for assessing sIOs’ response, we treated them with various cisplatin concentrations and evaluated the results using bright-field microscopy (magnification: ×100). Dead organoids appeared darker in contrast to those treated with the vehicle. Darkening of the dead organoids has also reported in ^18^. Based on our evaluation, we selected 10 µM as the optimal concentration for examining sIOs’ response to cisplatin in the viability assessment experiment.

**Extended Data Figure 3:**
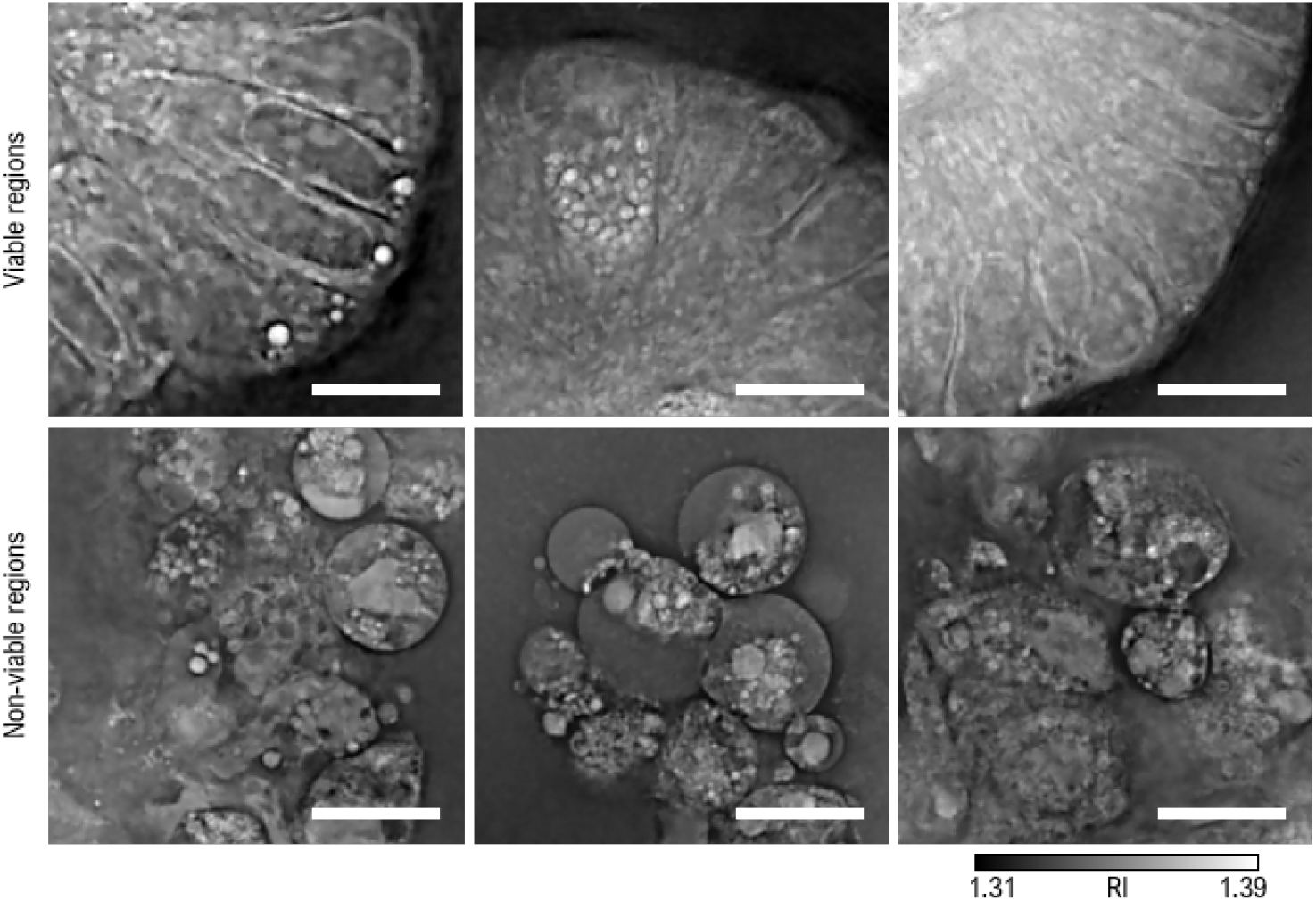
Variability in RI distribution of sIOs based on cell viability. Non-viable regions exhibit a low RI and lose their integrity compared to viable ones. Cells in the viable region display clear nuclei and maintain their polarity. Scale bar = 10 µm.

